# Phenotypic plasticity and genetic differentiation drive troglomorphic character development in European cave loach

**DOI:** 10.1101/2023.02.22.529532

**Authors:** Jasminca Behrmann-Godel, Samuel Roch, Alexander Böhm, Jolle Jolles, Alexander Brinker

## Abstract

Using a cross-fostering experiment, we provide evidence for the contribution of both genetic differentiation and phenotypic plasticity to troglomorphic character development in the recently discovered cave form of *Barbatula barbatul*a, an evolutionarily young lineage and first cavefish described in Europe, the northernmost record. We established reproducing populations of cave- and surface-dwelling loaches to produce cave, surface, and hybrid offspring and reared the F1 fish in a common garden setting in total darkness (DD) to simulate cave conditions as well as under the natural photoperiod (DL). We observed significant differences in the occurrence and extent of typical troglomorphic target characters among the offspring groups. Regardless of rearing conditions, cave fish exhibited smaller eyes, lighter body coloration, longer barbels, and larger olfactory epithelium than seen in surface fish. Hybrids in both rearing conditions generally showed an intermediate level of these traits. Surface and hybrid DD fish differed from the DL groups, resembling the cave fish phenotype in several traits, including eye size and body pigmentation. In contrast, cave and hybrid DL fish groups resembled surface fish phenotypes. Results confirmed that troglomorphic traits arise from heritable genetic differentiation of cave from surface forms and that phenotypic plasticity contributes to the process of adaptation to novel light conditions.

## Introduction

Originating from surface ancestors, cave-dwelling organisms typically evolve specific troglomorphic adaptations as a consequence of the strong selective pressures associated with living in darkness. The adaptations include the reduction and even complete loss of eyes and pigmentation and elongation of appendages associated with tactile and olfactory senses^1–3^. Stable and predictable conditions, particularly the absence of light, clearly indicate the direction of selection^4^. Thus cave organisms may offer a unique opportunity to better understand evolutionary principles.

Most known cave fishes occur in tropical and subtropical regions^5^. Species such as the Somalian desert cavefish *Phreatichthys andruzzii* have been isolated in subterranean environments over millions of years and lack epigean descendants or closely related surface species^6,7^. Others, like the Mexican cave tetra *Astyanax mexicanus* and fish of the genus Sinocyclocheilus in China have repeatedly colonized subterranean systems, with some cave populations being of more recent origin^8–11^. Few cave fish species are known in temperate regions^12–14^, one of these being the Aach cave loach, representing the first, and currently only known, cave fish in Europe. Behrmann-Godel et al.^15^ established that stone loach *Barbatula barbatula* entered the Danube-Aach subterranean system in the most recent post-glacial period. Since the region was glaciated until the end of the Late Pleistocene (Würm) glacial period^16,17^, loach would have colonized the underground system within the past 20,000 years. Specimens captured in the underground system show typical troglomorphic characters: reduction in eye size, pale body pigmentation, enlarged olfactory epithelium, and longer barbels compared to the surface form that occurs in the connected waters of the Danube and Radolfzeller Aach Rivers^15^. Genetic comparisons have revealed that the cave-dwelling loach is more closely related to surface stone loach present in the River Danube than that from the Radolfzeller Aach^15^.

Surface fish entering a subterranean environment will encounter immediate, and likely severe, selection pressures that will eventually result in troglomorphic characteristics and associated genetic divergence from its ancestral population^2,4^. In addition, the novel environment *per se* might directly induce alterations in morphology, physiology, and behaviour. Such changes, which uncouple phenotype from genotype, reflect phenotypic plasticity^18^, an individual response to novel environmental conditions. Alterations in trait expression are typically within a genotype-specific reaction norm^19^. This reaction norm might provide a survival advantage to individuals tolerating the new environment^20^, possibly resulting in an initial step that is subsequently reinforced by genetic change and natural selection^21^. However, over the long term, individual plasticity might be selected *against* in organisms trapped in an environment in which conditions are stable, such as permanent darkness^22^. Rohner et al.^23^ provided evidence that variation in orbit size in eyeless cave-dwelling *A. mexicanus* is lower than that seen in surface forms of the fish. In a permanently dark environment, vision provides no benefit. Development of the vision system is costly^24^, and there may be a selective advantage in eye regression, with the complete loss of eyes becoming heritable and finally genetically fixed^22^.

A fundamental question when working with an evolutionarily young species like the European cave-dwelling *B. barbatula*, with surface forms present in the same aquatic system, is how far the organism has diverged from its epigean ancestors. In such a young lineage, morphological, behavioural, physiological, and genetic differences between epigean and cave forms can be analysed and the adaptation process studied^2^. It might also be possible to interbreed surface and cave fish. Such crosses provide a unique opportunity to study the restoration of typical characters and reveal the genetic basis of the process of troglomorphic trait evolution^10^.

With the goal of obtaining fundamental insight into the environmental adaptation process in cave-dwelling *B. barbatula*, we raised F1 offspring in a common garden setting that reflected either cave conditions (complete absence of light), or surface conditions (natural dark/light cycle) and compared their eye size, lens quality, barbel length, size of olfactory epithelium, and the level of body pigmentation. We hypothesize that, if genetic differentiation between surface and cave forms has been established with heritable troglomorphic characters, cave fish would express a unique phenotype regardless of light conditions. Hybrids should possess intermediate character states for a single trait, a character state that resembles one of the parental forms, or some combination of the two, depending on effects of genetic variation, dominance, and epistasis. We also hypothesize that, if phenotypic plasticity contributes to adaptation to a cave habitat, there should be pronounced phenotypic variation depending on light conditions similar to differences typically seen between cave and surface fish.

## Material and Methods

### Specimen source and maintenance

Fifty-four cave loach were captured from the Danube-Aach cave system by a professional cave diver via hand netting (Fig.1a and b; Supplementary methods). From one to 11 fish were captured on each of twenty-one dives from 2015 through 2018 (Fig. 1b). At the surface, the surviving specimens were carefully transferred to an aerated darkened bottle and immediately transported to the laboratory. Sixty-one surface stone loach were caught in spring 2016 from the Danube and the Radolfzeller Aach rivers by electrofishing (Fig. 1a) as described^15^.

**Fig. 1.**
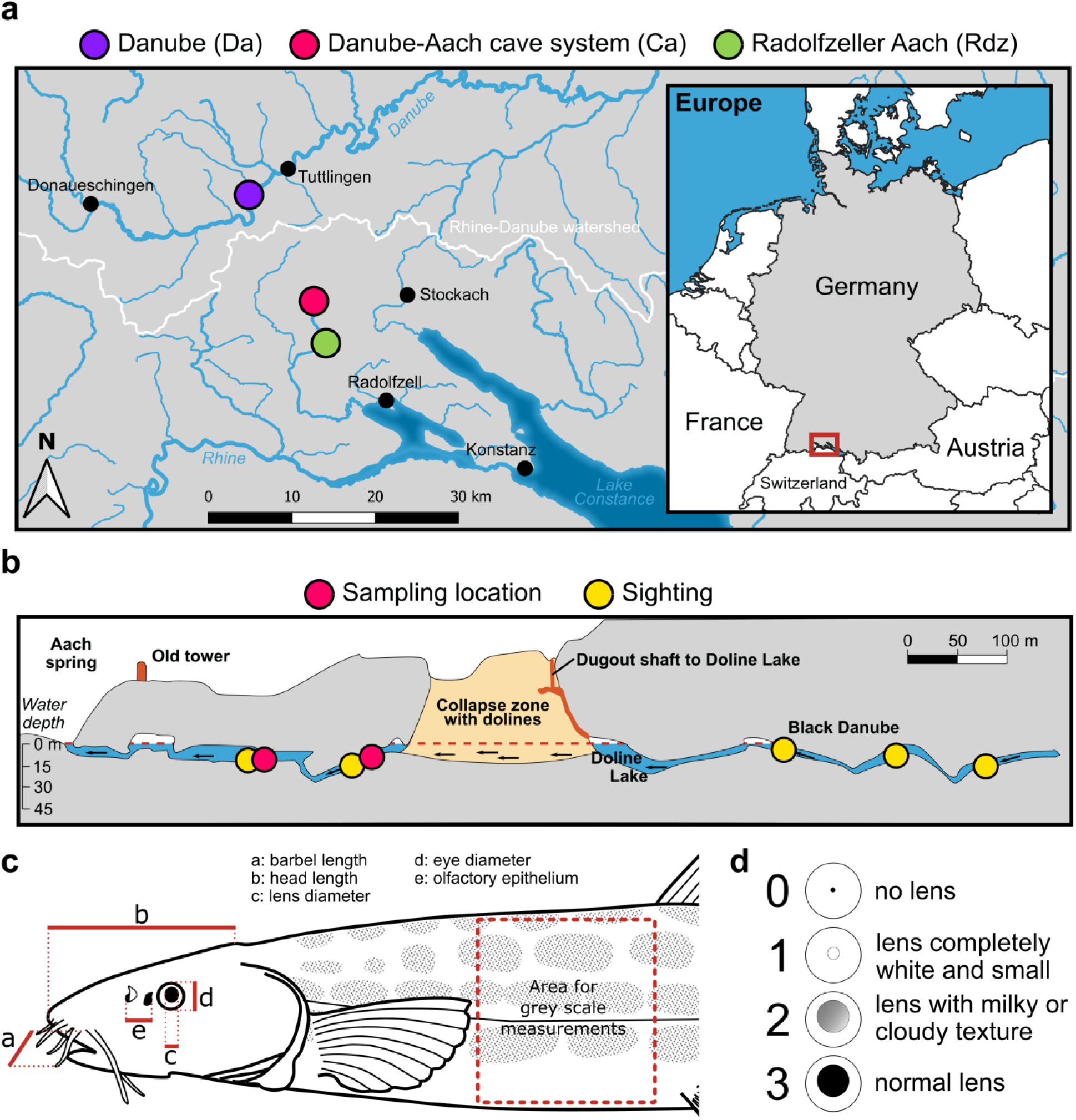
Study site and measured phenotypic traits. **(a)** Sampling locations for cave and surface loach; precise location is shown in Behrmann-Godel et al. (2017). **(b)** Schematic of the Danube-Aach cave system near the Aach spring. Entry is from the Aach Spring or via the dugout shaft leading to Doline Lake. Dots represent cave loach sightings during 2015– 2022. Arrows indicate water flow from Danube to Aach spring. **(c)** Selected morphometric parameters to assess troglomorphic characters. **(d)** Definition of lens quality classes.

Twenty cave specimens were separated into size-based groups of 9, 8, and 3 and stocked into 60 l tanks (60 × 45 × 34 cm) in complete darkness. A single white light of low amperage was used when feeding and cleaning the aquaria. Twenty loach from the Danube and 10 from the Radolfzeller Aach were held in similar tanks under the natural photoperiod. All tanks had a layer of sand and gravel substrate, with stones and PVC pipe added for shelter and an air stone for aeration. A constant flow-through of filtered (100 µm) and aerated water taken directly from Lake Constance at a depth of ∼20 m was provided. Water temperature was not altered from lake temperature until breeding. Fish were fed dry food, frozen chironomids, and live daphnia from the institute’s breeding ponds *ad libitum*.

### Cross-fostering to produce F1 cave, surface, and hybrid fish

We initiated controlled pairwise breeding for common garden experiments in spring 2019. The extended period from collecting fish until breeding guaranteed adequate habituation, acclimation to laboratory conditions, and maturation. The offspring exposed in the common garden experiments were progeny of the wild-caught fish with the exception of two male F1 fish from an earlier breeding of our captured population (1 cavefish, 1 Rdz), as we lacked a sufficient number of male broodstock (Supplementary Methods). Twenty pairs consisting of a single male and female of specified origin (cave, Danube, Radolfzeller Aach) were placed in individual circular 20 l tanks (Table 1; Supplementary Methods).

**Table 1.** Breeding of fish for common garden experiments. Origin of male and female parents, number of egg clutches produced per pair, and number of hatchlings per clutch. The final two columns show the number of fish used for analysis of morphology. Da = Danube, Rdz = Radolfzeller Aach, Ca = cave, Hy = hybrid, F1 = first laboratory-bred generation. DD = completely darkness, DL = natural day/night cycle. NA = successful hatching, number of hatchlings not analysed.

| ID family | Male origin | Female origin | No. of clutches | Hatchlings Per clutch | DD [no. fish] | DL [no. fish] |
| --- | --- | --- | --- | --- | --- | --- |
| Do1 | Da | Da | 1 | 150 | 17 | 10 |
| Do2 | Da | Da F1 | 1 | 300 | 4 | 15 |
| Do3 | Da | Da | 0 | - | - | - |
| Do4 | Da | Da | 2 | 300 | 4 | 0 |
| RdZ1 | Rdz | Rdz | 1 | 120 | 7 | 0 |
| RdZ2 | Rdz | Rdz | 0 | - | - | - |
| RdZ3 | Rdz | Rdz | 1 | 100 | 9 | 12 |
| RdZ4 | Rdz | Rdz | 2 | 200/NA | 9 | 13 |
| C1 | Ca | Ca | 0 | - | - | - |
| C2 | Ca | Ca | 2 | 110/NA | 0 | 0 |
| C3 | Ca F1 | Ca | 2 | 110/100 | 12 | 13 |
| C4 | Ca | Ca | 0 | - | - | - |
| C5 | Ca | Ca | 1 | NA | - | - |
| C6 | Ca | Ca | 1 | 80 | 13 | 12 |
| Hy 1 | Rdz F1 | Ca | 2 | NA/150 | 21 | 21 |
| Hy 2 | Rdz | Ca | 2 | 20/10 | - | - |
| Hy 3 | Da | Ca | 1 | 20 | - | - |

### Common garden experiments

All eggs used for the common garden experiments were produced from 10 April through 7 June 2019. Each egg clutch (50–500) was separated into two batches immediately after detection, usually in early morning, and transferred into one of two rooms equipped with circular 20 l tanks with water flow-through from Lake Constance and an air stone added for aeration. One room simulated cave conditions and lacked any light source (DD). During feeding and maintenance, a red LED light strip provided a single temporary light source. The second room had windows plus an artificial light regime following the natural seasonal photoperiod (DL). Other conditions, including water temperature (14°C) and feeding regime, were identical.

Breeding was challenging. We experienced high egg mortality during the first days post-fertilization, probably due to fungal infections, and survival from egg to late juvenile stage was not always successful, sometimes resulting in only a small number of progeny per pair. Nevertheless, we obtained 19 spawning events from 13 fish pairs, with six pairs producing more than one fertilized egg clutch (Table 1). Ten to 300 (Table 1) juveniles hatched ∼10 days post-fertilization and started feeding approximately five days after hatching. Fry were fed freshly hatched *artemia nauplii* and crushed flake commercial food (Tetra) *ad libitum*. Tanks were cleaned every three days. Food particle size was increased as fish grew, and live *Daphnia* and frozen bloodworms were added as appropriate (Supplementary Methods).

The common garden experiments were terminated on 16 and 17 August 2019 for the DL and DD treatments, respectively, when all fish were at least two months old. Fish were netted from the 20 l holding tanks, anesthetized with MS-222 (0.1 g/l), placed individually in a small Petri dish on the left side, and photographed under identical camera and light settings to measure total body length and for grey-scale analysis of body pigmentation (Fig. 1c). Fish were then moved to individual holding tanks where they were kept for behaviour observations (data not included) after which they were euthanized with MS-222 (3 g/l), transferred to formalin (4%), and stored at 4 °C until analysis.

Ninety-six fish were used to assess morphology; 25 each of cave, Danube, and Radolfzeller Aach origin and 21 hybrids (Table 1). To avoid familial effects, we attempted to use a similar number of fish from each parent pair. If enough fish were available, the first clutch of a pair was used. When the number of fish from the first clutch was insufficient, those from the second clutch were included (Table 1).

### Data processing

Photos of anaesthetized fish were examined to compare pigmentation of fish with grey-scale analysis. A photo was opened in ImageJ^25^ and a representative rectangular area between the pelvic fin and the beginning of the dorsal fin was selected (Fig. 1c). The grey-scale measurement tool by ImageJ provides a number from 0 (black) to 254,000 (white) representing pigmentation of the individual fish.

All morphometric measurements were taken from photographs of formalin-fixed fish. Specimens were photographed with a binocular microscope from the left and from above using the ImageJ program (Fig. 1c; Supplementary Fig S1). Lateral images were used to measure the diameter of the eye and lens and the length of the second barbel (Supplementary Fig S1a). Head length was measured from the tip of the snout to the end of the skull bone (Supplementary Fig S1b). The olfactory epithelium in *B. barbatula* is an oval area located between the inflow/anterior and the outflow/posterior nostrils on each side of the fish snout^26^. The size (longest axis) of the olfactory epithelium was measured as distance between the anterior and the posterior nostril on one side of the fish (Fig. 1c; supplementary Fig. S1b). All length measurements were done to the nearest µm.

Using the lateral view, lenses were categorized into one of four classes to indicate quality. Class 3 (normal): lens clear with a circular shape of approximately half the diameter of the eye; Class 2: lens milky or cloudy, diameter 80–100% of class 3 lenses; Class 1: lens < 80% the size of Class 3, milky, or completely white; and Class 0, no lens detected (Fig. 1d).

### Data analysis

General linear models were used to analyse eye diameter, lens diameter, barbel length, olfactory epithelium size, lens quality, and grey-scale using the formula

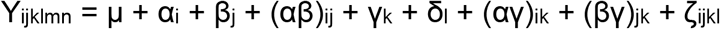

where Y_ijklmn_ is morphometric or colour trait; µ is the overall mean, α_i_ denotes origin, β_j_ is treatment (DD or DL), (αβ)_ij_ is interaction, γ_k_ is total length, δ_l_ is age (days), (αγ)_ik_ is the interaction between origin and length, (βγ)_jk_ is the interaction between treatment and length, and ζ_ijkl_ is the random residual error. Model assumptions were tested using normal quantile plots of residuals. *Post hoc* comparisons were made with Tukey’s HSD test (origin), Student-t test (treatment), or by constructing contrasts (origin *vs*. treatment). To compare variability in eye and lens diameter among origins, we calculated the eye diameter and barbel length relative to body length and the lens diameter relative to eye diameter. Subsequently, we determined the coefficient of variation (CV) for each origin, and tested pairwise differences by sequential Bonferroni-corrected Levene tests. All statistics were run on JMP Pro 16.2.0 (64-bit, SAS Institute). Significance was set at p < 0.05.

### Ethics

The study was carried out in accordance with the Protection of Animals Act Germany. All the animals were maintained and handled according to the Institutional Animal Care guidelines of the University of Konstanz. Animal holding was approved by the regional council Freiburg (reference number 35-9185.64/1.1). Animal experimentation was approved by the regional council Freiburg (reference number: 35-9185.81/G-19/30). Animal collection was approved by the regional council Tübingen (reference number 33-4/9220.51-3). Approval by a review board institution or ethics committee for fishing wild loach was not necessary, as fish were caught by permission of the local fisheries administration (Fisheries Administration RP Tübingen).

## Results

### Vision system

In general, eye and lens size as well as lens quality differed extensively between cave and surface fish.

Lens quality was strongly linked to fish parental origin, whereas treatment, total length, and age showed no significant effect (Table 2). More than 95% of all cave fish progeny showed impaired lens quality with >60% falling into classes 0 and 1, meaning lenses were either non-existent or completely white (Fig. 2). In contrast, hybrid and surface fish offspring nearly all had normal lens quality, and <3% of hybrid fish exhibited category 2, lenses with milky or cloudy texture (Fig. 2).

**Table 2.**
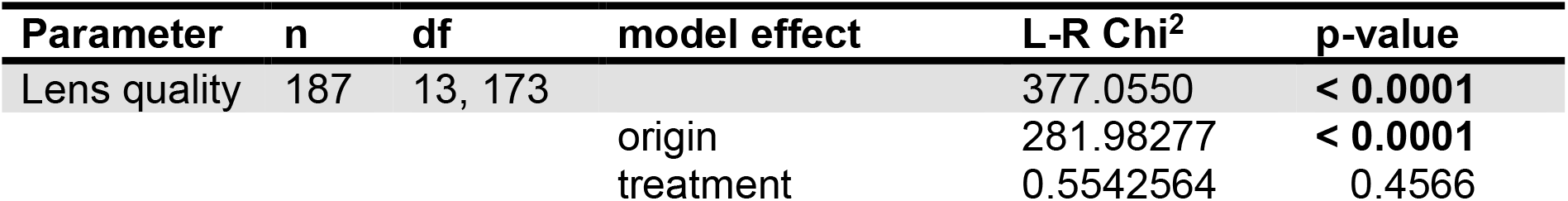
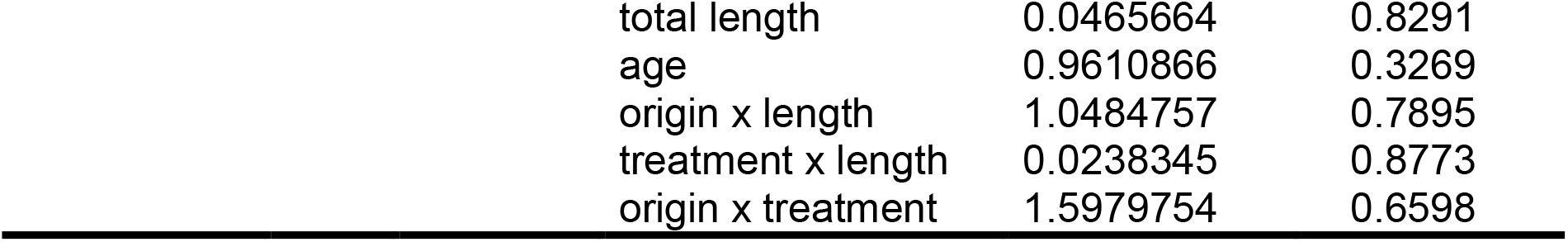
Results of the general linear model analysis of loach lens quality. n = number of observations. df = degrees of freedom. Significant differences (p < 0.05) are in bold.

**Fig. 2.**
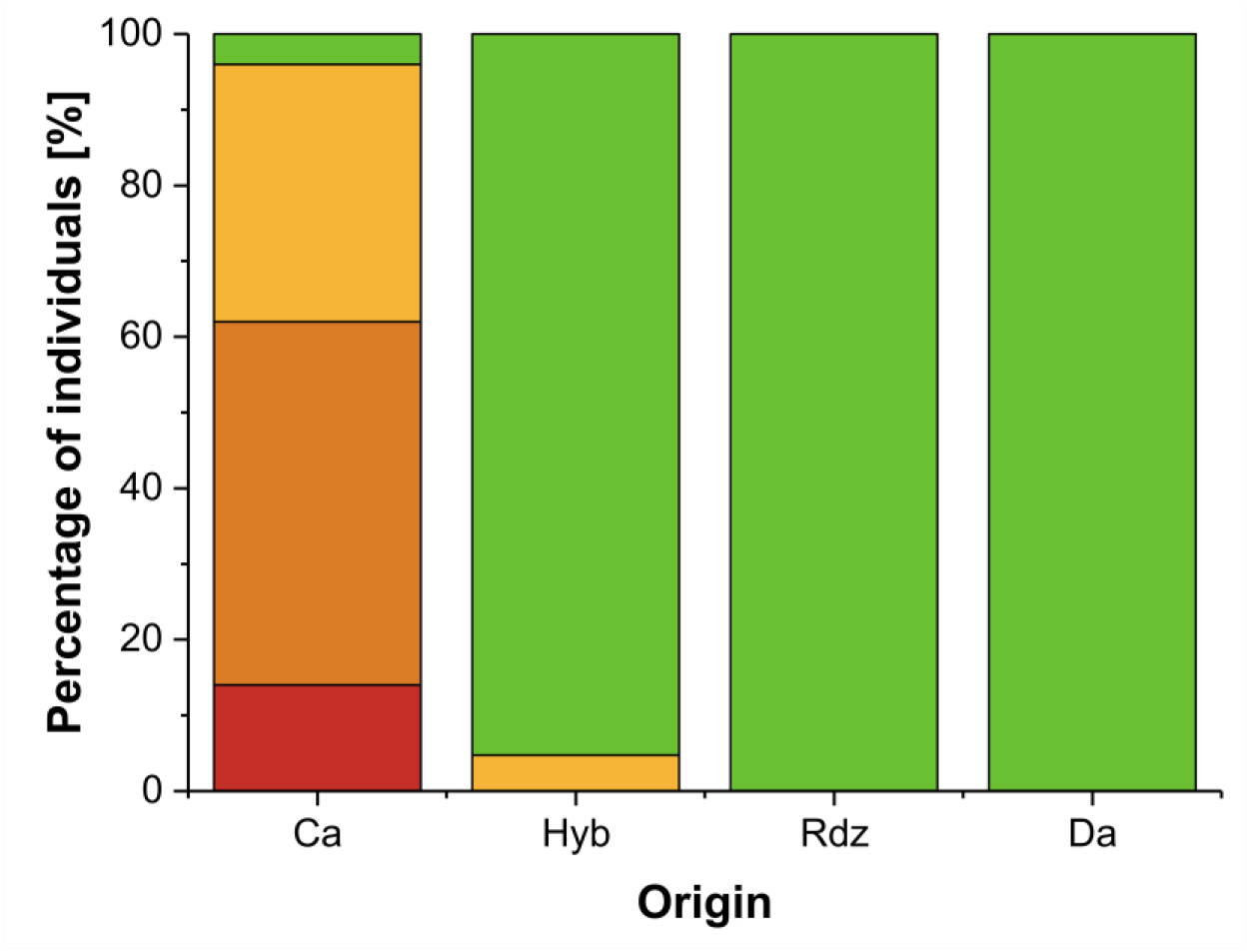
Lens quality. Proportion of loach lens classes with respect to parent origin (Ca = Cave, Rdz = Radolfzeller Aach, Da = Danube, Hyb = hybrids). Red: class 0; orange: class 1; yellow, class 2; green, class 3 (Fig. 1D). For GLM results, see Table 2.

Differences in allometrically-corrected eye and lens diameter of the F1 fish was best explained by parent origin (Table 3). All cave fish offspring had significantly smaller eyes (1.0 ± 0.2 mm) than offspring of surface fish (Radolfzeller Aach = 1.4 ± 0.2 mm, Danube = 1.6 ± 0.2 mm) (Fig. 3). Hybrid eye diameter was intermediate between that of cave and surface fish (1.3 ± 0.1 mm). Eye diameter of Radolfzeller Aach fish was lower than that of the Danube fish (Fig. 3). Similarly, lens diameter was significantly lower in cave fish (0.2 ± 0.1 mm) compared to surface fish (Radolfzeller Aach = 0.6 ± 0.1 mm, Danube = 0.6 ± 0.1 mm), with hybrids having lenses of intermediate size (0.4 ± 0.04 mm; Fig. 3). Total length and, to a lesser extent, fish age were also significantly related to eye and lens diameter, with longer and older fish exhibiting larger eyes and lens diameters (Fig. 3, Table 3). Finally, a small but significant effect (p = 0.0129) of treatment on eye diameter was observed (Table 3), with the fish of similar origin reared in permanent darkness having smaller eyes than those reared under the natural photoperiod (Fig. 3). We did not find such an effect for lens size (Table 3).

**Table 3.**
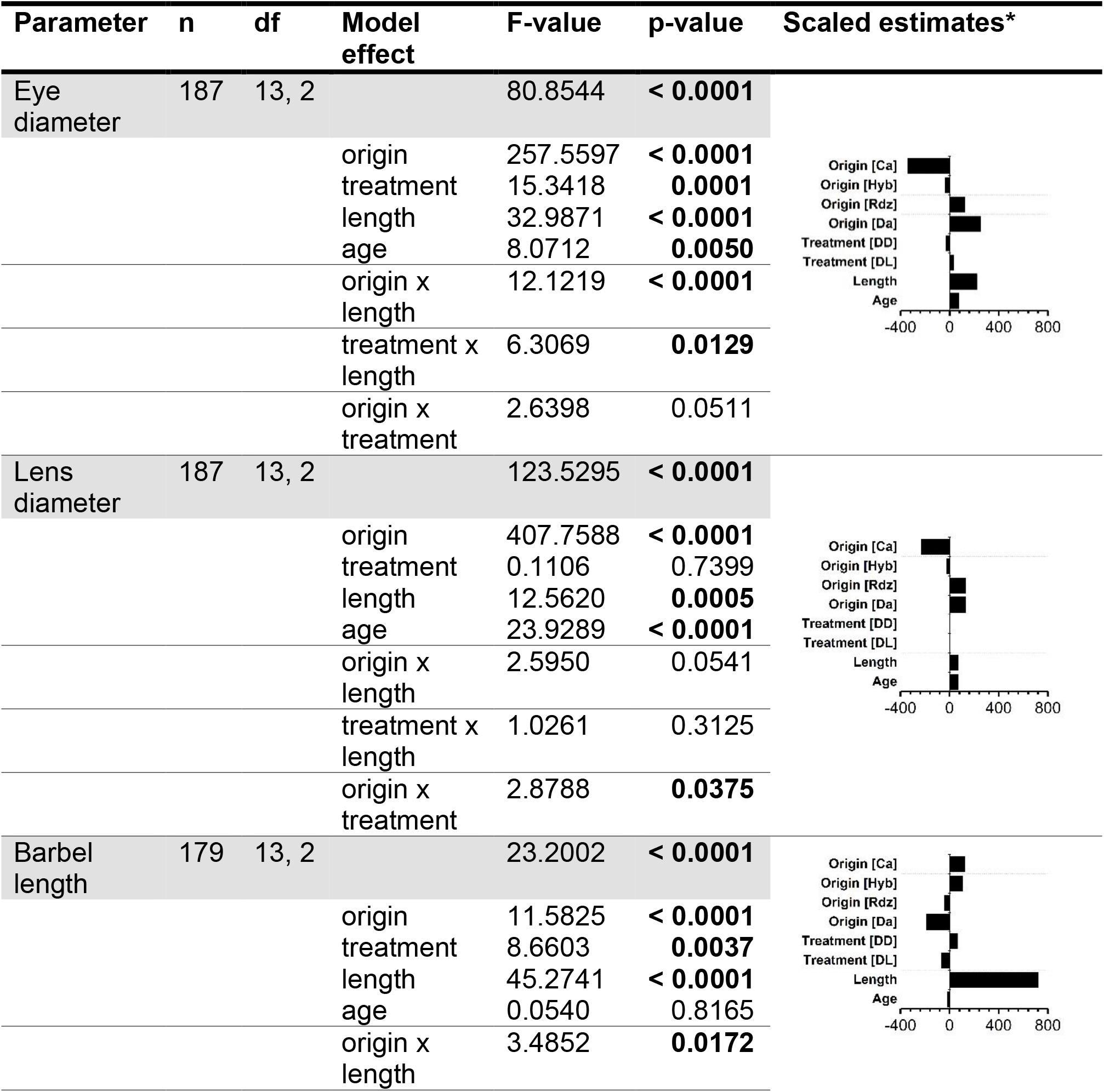
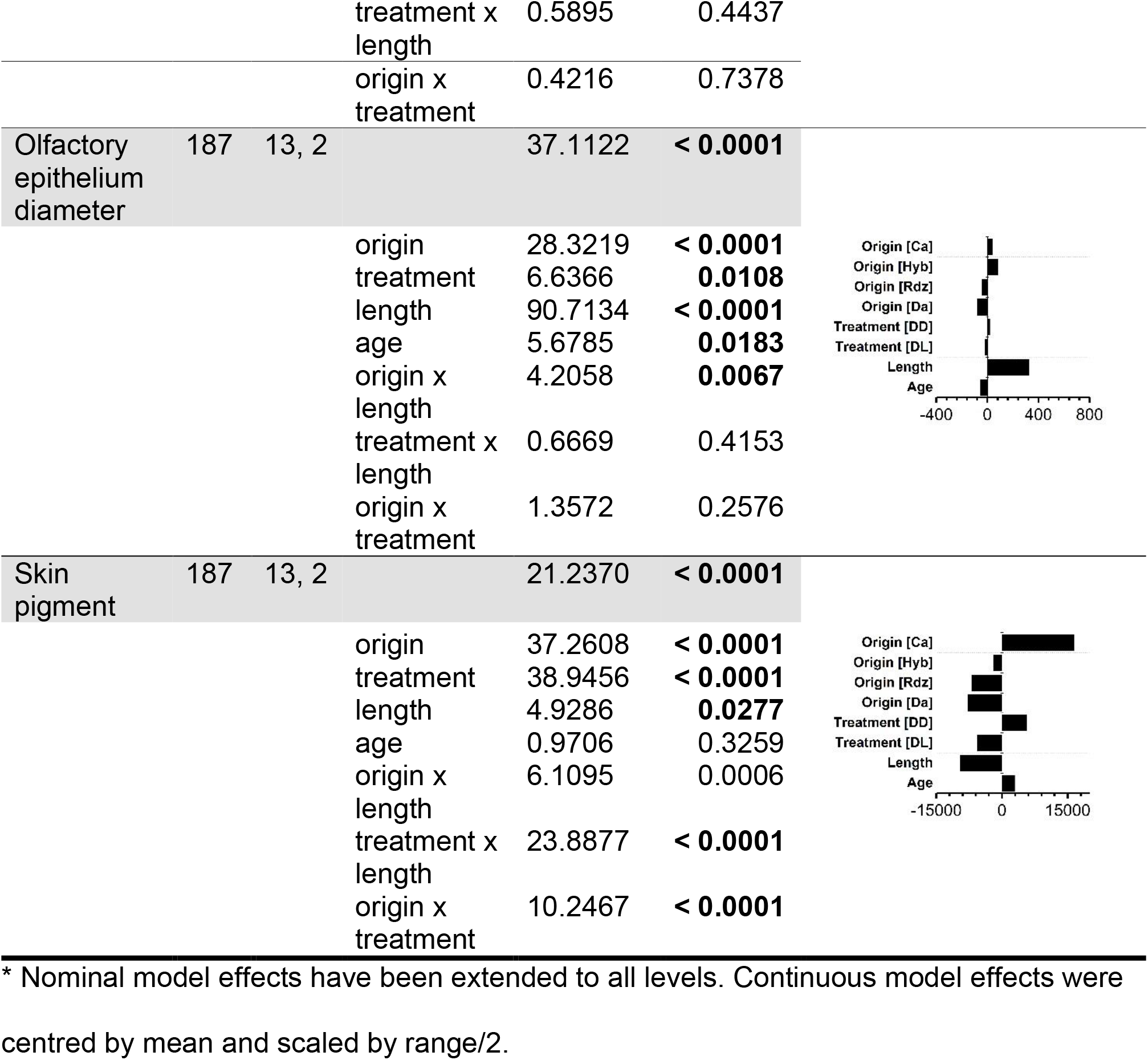
Results of general linear modelling of selected morphometric parameters of loach. n = number of observations. df = degrees of freedom. Da = Danube, Rdz = Radolfzeller Aach, Ca = Cave, Hyb = hybrids. DD = darkness, DL = natural day/night cycle. Significantly different values (p < 0.05) are highlighted in bold.

| Parameter | n | df | Model effect | F-value | p-value | Scaled estimates* |
| --- | --- | --- | --- | --- | --- | --- |
| Eye diameter | 187 | 13, 2 |  | 80.8544 | **< 0.0001** |  |
|  |  |  | origin | 257.5597 | **< 0.0001** |  |
|  |  |  | treatment | 15.3418 | **0.0001** |  |
|  |  |  | length | 32.9871 | **< 0.0001** |  |
|  |  |  | age | 8.0712 | **0.0050** |  |
|  |  |  | origin x length | 12.1219 | **< 0.0001** |  |
|  |  |  | treatment x length | 6.3069 | **0.0129** |  |
|  |  |  | origin x treatment | 2.6398 | 0.0511 |  |
| Lens diameter | 187 | 13, 2 |  | 123.5295 | **< 0.0001** |  |
|  |  |  | origin | 407.7588 | **< 0.0001** |  |
|  |  |  | treatment | 0.1106 | 0.7399 |  |
|  |  |  | length | 12.5620 | **0.0005** |  |
|  |  |  | age | 23.9289 | **< 0.0001** |  |
|  |  |  | origin x length | 2.5950 | 0.0541 |  |
|  |  |  | treatment x length | 1.0261 | 0.3125 |  |
|  |  |  | origin x treatment | 2.8788 | **0.0375** |  |
| Barbel length | 179 | 13, 2 |  | 23.2002 | **< 0.0001** |  |
|  |  |  | origin | 11.5825 | **< 0.0001** |  |
|  |  |  | treatment | 8.6603 | **0.0037** |  |
|  |  |  | length | 45.2741 | **< 0.0001** |  |
|  |  |  | age | 0.0540 | 0.8165 |  |
|  |  |  | origin x length | 3.4852 | **0.0172** |  |

|  |  |  |  |  |  |
| --- | --- | --- | --- | --- | --- |
|  |  |  | treatment x length | 0.5895 | 0.4437 |
|  |  |  | origin x treatment | 0.4216 | 0.7378 |
| Olfactory epithelium diameter | 187 | 13, 2 |  | 37.1122 | **< 0.0001** |
|  |  |  | origin | 28.3219 | **< 0.0001** |
|  |  |  | treatment | 6.6366 | **0.0108** |
|  |  |  | length | 90.7134 | **< 0.0001** |
|  |  |  | age | 5.6785 | **0.0183** |
|  |  |  | origin x length | 4.2058 | **0.0067** |
|  |  |  | treatment x length | 0.6669 | 0.4153 |
|  |  |  | origin x treatment | 1.3572 | 0.2576 |
| Skin pigment | 187 | 13, 2 |  | 21.2370 | **< 0.0001** |
|  |  |  | origin | 37.2608 | **< 0.0001** |
|  |  |  | treatment | 38.9456 | **< 0.0001** |
|  |  |  | length | 4.9286 | **0.0277** |
|  |  |  | age | 0.9706 | 0.3259 |
|  |  |  | origin x length | 6.1095 | 0.0006 |
|  |  |  | treatment x length | 23.8877 | **< 0.0001** |
|  |  |  | origin x treatment | 10.2467 | **< 0.0001** |

\* Nominal model effects have been extended to all levels. Continuous model effects were centred by mean and scaled by range/2.

**Fig. 3.**
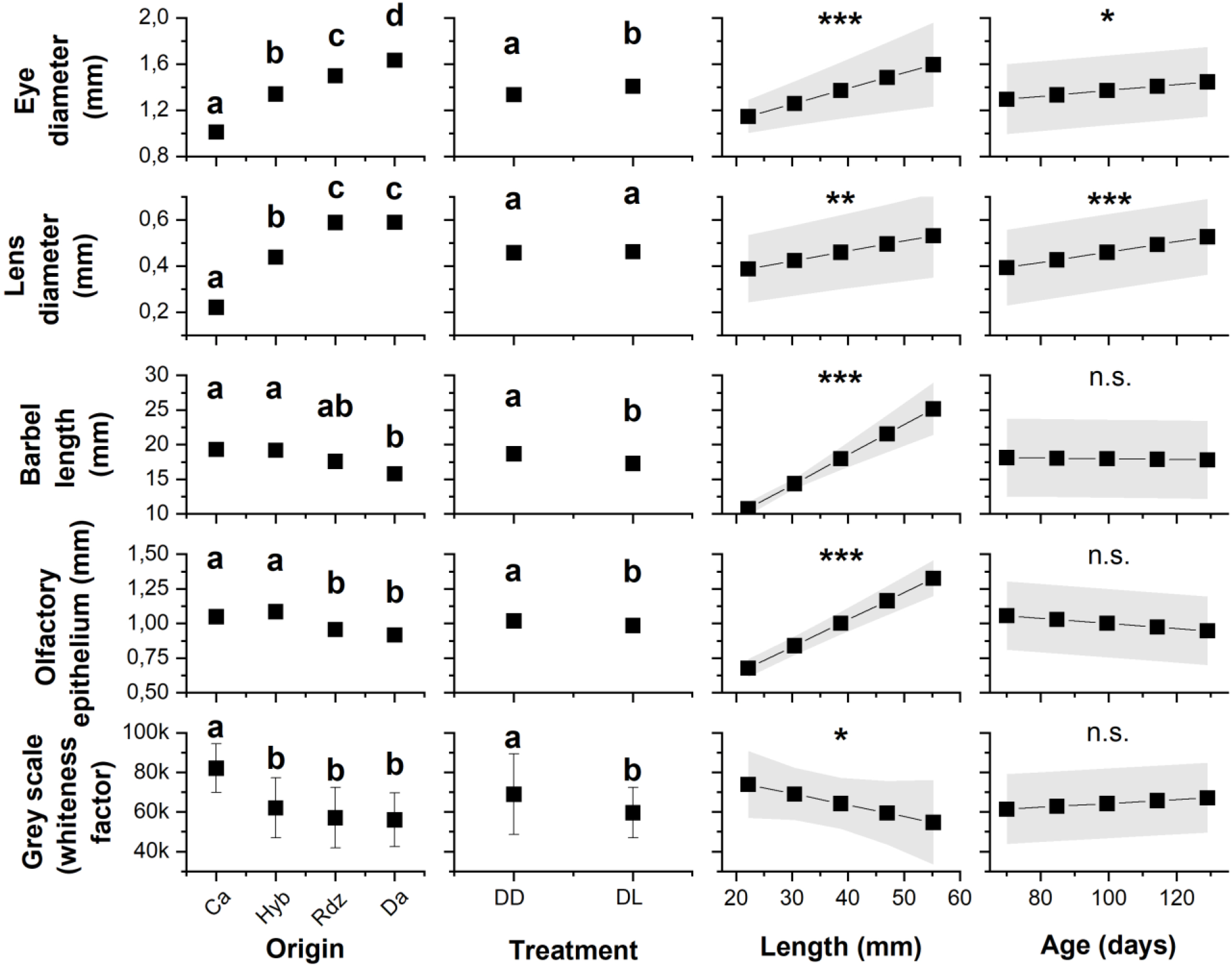
Results of generalized linear modelling to analyse selected characters. Shown are grand marginal means of selected parameters of loach relative to parent origin (Ca = Cave, Hyb = hybrids, Rdz = Radolfzeller Aach, Da = Danube) and treatment: total darkness (DD) or natural photoperiod (DL). Error bars and grey bands indicate standard deviation. For categorical data, significant differences are indicated by lowercase letters (Tukey HSD post hoc test). For continuous data, significant differences are indicated by asterisks (* p < 0.01, ** p < 0.001, *** p < 0.0001); for detail see Table 3.

The coefficient of variation in eye diameter was significantly greater in cave fish (CV = 23.18%) compared to surface (Radolfzeller Aach: CV = 9.27%, Danube: CV = 9.25%) and hybrid fish (CV = 5.73%) (Fig. 4, Table 4). While there was no significant difference in eye diameter variation between the two surface fish populations, hybrids exhibited significantly lower variation in eye diameter compared to other origins (Table 4). A similar pattern was found for variation in lens diameter: large in cave fish with CV = 47.44%, with some having no lens and others with lens sizes comparable to those of surface fish (Fig. 4; Table 4). In contrast, there was little variation in lens size among the surface (Radolfzeller Aach: CV = 6.18%, Danube: CV = 5.56%) and hybrid fish (CV = 6.30%), and no significant differences between them (Table 4).

**Table 4.** Results of the pairwise comparison of variation in eye and lens size of loach relative to parent origin. Pairwise statistical testing via contrasts used a Bonferroni-corrected Levene test; significant differences (p < 0.05) are highlighted in bold. Ca = Cave, Hyb = hybrids, Rdz = Radolfzeller Aach, Da = Danube.

| Contrasts<br>pairs tested | Eye size<br>p-value | Lens size<br>p-value |
| --- | --- | --- |
| Ca vs. Hyb | **<0.001** | **<0.001** |
| Ca vs. Da | **0.007** | **<0.001** |
| Ca vs. Rdz | **0.003** | **<0.001** |
| Da vs. Rdz | 0.818 | 0.194 |
| Da vs. Hyb | **0.004** | 0.580 |
| Rdz vs. Hyb | **0.003** | 0.422 |

**Fig. 4.**
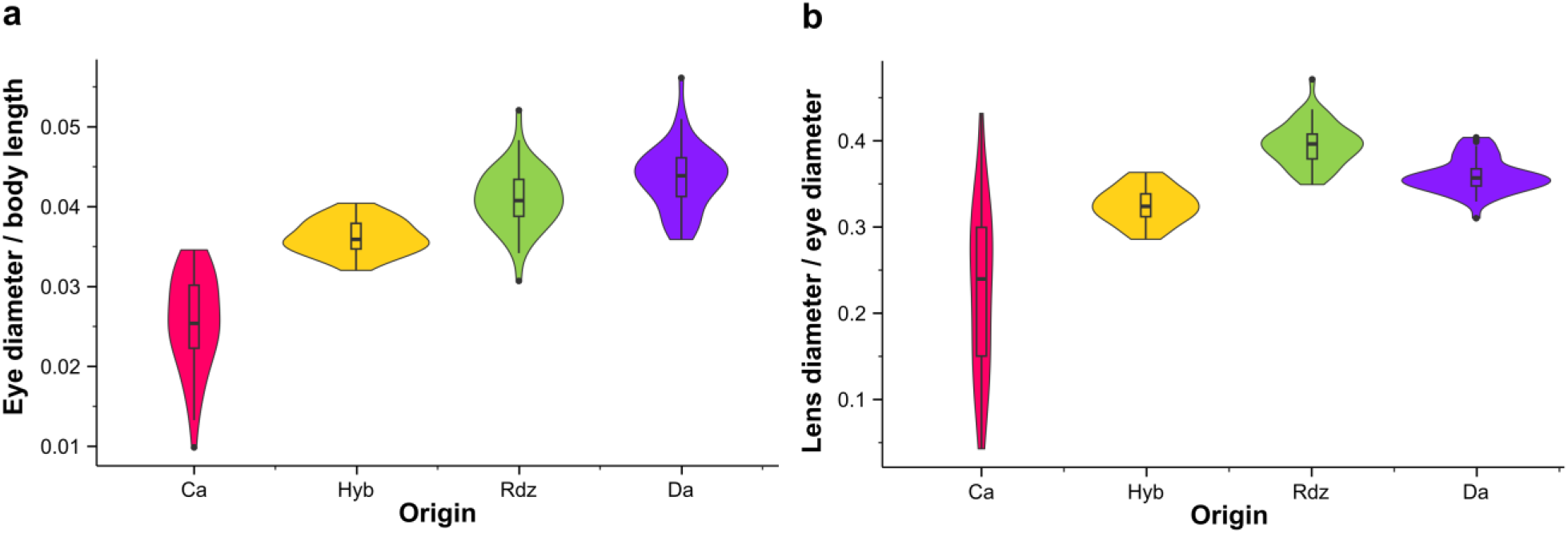
Differences in variation of eye and lens diameter. Shown are combined box and violin plots to illustrate variation in eye (**a**) and lens (**b**) size of loach with respect to parental origin (Ca = Cave, Hyb = hybrids, Rdz = Radolfzeller Aach, Da = Danube).

All cave fish F1 offspring exhibited eyes retracted into the head and often not visible from the overhead camera angle, similar to wild fish collected in the cave system (supplementary Fig. S1b). We did not observe the lateral asymmetry in eye development seen in three cave fish collected from the wild (supplementary Fig. S2a–c).

### Barbel length

The best predictor of second barbel length was fish total length (Table 3), with longer fish having longer barbels (Fig. 3). Origin of parent stock and treatment showed significant effects (Table 3). Cave fish (2.0 ± 0.6 mm), and hybrids (1.8 ± 0.2 mm) had the longest barbels, followed by fish from the Radolfzeller Aach (1.6 ± 0.4 mm) and finally the Danube (1.5 ± 0.2 mm) (Fig. 3). Regardless of parent stock, fish reared in darkness developed significantly longer second barbels than those in the natural photoperiod (Fig. 3).

### Olfactory epithelium

Olfactory epithelium size was significantly positively correlated with fish total length (Table 3), (Fig. 3) and was significantly longer in cave fish (1.1 ± 0.2 mm) and hybrids (1.0 ± 0.1 mm) than in surface fish regardless of treatment (Radolfzeller Aach = 0.9 ± 0.1 mm, Danube = 0.9 ± 0.1 mm) (Fig. 3). Treatment showed a weak but significant effect in fish of similar origin, with those reared in darkness exhibiting larger olfactory epithelium than those under natural light/dark conditions (Fig. 3).

### Pigmentation

Fish parent origin and treatment were both good predictors of degree of pigmentation (Table 3). Cave fish had lower pigmentation than hybrids and surface fish, (Fig. 3) with this effect stronger expressed in darkness, but the difference was not significant. Fish raised in darkness were, on average, less pigmented than fish raised under the natural photoperiod (Fig. 3). Total length was significantly related to fish pigmentation (Table 3), with smaller fish being less pigmented than larger fish (Fig. 3). Age had no significant effect (Table 3).

## Discussion

We established the first laboratory population of the recently-discovered European cave loach and crossed wild cave fish with surface populations. The European cave loach exhibited heritable troglomorphic traits similar to those of other cave fish species ^2,10^, confirming that, despite being an evolutionarily young lineage^15^, they have substantially adapted to cave conditions. This was especially evident in the expression of regressive traits, in particular the reduction of the eye and lens diameter, the loss of functional lenses, and the reduction in pigmentation as well as, to a lesser extent, constructive traits including the elongation of barbels and increase of the size of the olfactory epithelium.

For the population of cave fish studied, the adaptive process has not resulted in complete eye loss, although eye regression can be clearly observed. This is also manifested in the wide range of expression of other troglomorphic characters in cave loach (Behrmann-Godel *et al*.^15^). The phenomenon provides a unique opportunity to distinguish plastic responses to acute environment change from heritable genetic differentiation developed over a long period of adaptation.

The lack of light is the most obvious difference in cave and surface environments. Vision is a primary sense in most fish species, and a key factor in the entrainment of circadian rhythms, orientation, navigation, detection of predators, foraging, and mating^27^. The development and maintenance of eyes is energetically costly, hence their loss is advantageous when no longer functional^24,28^. In the present study, all laboratory-reared cave loach offspring exhibited limited development of the vision system. The eyes and lenses of hybrids were intermediate in size of their parental forms, while, with the exception of a single specimen, the quality of the lens was indistinguishable that from surface fish, with almost all hybrids showing normal lenses. These findings indicate aspects of genes or gene function that underlie lens formation and function in cave fish that might already be fixed in the cave fish population. Whether these changes are based on coding or non-coding mutations or an alteration of the DNA methylation pattern in the genes associated with lens degeneration needs to be investigated. The high variation of eye size in cave loach might reflect its early evolutionary history or could indicate ongoing introgression of surface and cave fish. Future studies that include genetic and genomic analysis might help explain these findings.

The basis for eye loss in cave organisms is debated^29^ and may represent specific progressive steps, including epigenetic changes that deactivate expression of relevant genes, inactivating mutations, gene loss, and fixation of genetic variants^10,30,31^. Although several studies have analysed genes related to eye development in the eyeless *A. mexicanus* from the Pachon cave, no inactivating mutations have been found^32–36^. Nevertheless, a recent study did find numerous changes in DNA methylation patterns to be responsible for the regression of several eye development genes in the cave form of *A. mexicanus*^30^. It was the first to show that epigenetic modification can play a major role in eye degeneration during cave fish embryonic development. In addition to variability in the genes and pathways responsible for the selection of adaptive phenotypes in cave-dwelling organisms, other evolutionary processes might be involved and help explain strong troglomorphic effects observed in the F1 offspring of the present study^31^.

In our experiments, cave loach were consistently less pigmented than surface fish, irrespective of rearing conditions. This indicates trait regression, which is a typically observed change in cave fish^2^. However, several laboratory-bred cave loach not included in this experiment that were reared in the natural photoperiod gradually acquired darker pigmentation, reaching a plateau at about six months of age (data not shown). This reversibility in pigmentation reduction is found in a number of cave fish species and in the cave salamander *Proteus anguinus* ^37^. However, while some cave loach kept in natural light conditions developed a darker body over time, they never exhibited the typical pattern of dark blotches observed in surface stone loach, and reversibility of pigmentation was not observed in all cave fish offspring. Cross-breeding experiments in *A. mexicanus* showed pigment reduction to be a polygenic trait in which multiple genes, of which only a handful are known, have repeatedly been the target of mutation^36^. If pigmentation reduction in the cave loach is the effect of mutation of several genes, the high variation in character state observed might be the result of an inherited individual mutation pattern. Following this logic, there may be a mosaic of mutation patterns in our cave fish population, of which some combinations may result in more or less pigmentation in offspring as well as the potential for pigmentation reversal.

While it is widely accepted that eye regression in cave animals is ultimately the result of natural selection^38,39^, the mechanisms driving pigmentation loss in cavefishes is less clear. One oft-held opinion is that it is the result of neutral mutation and drift, as indicated by polarities in QTL mapping of melanophore number in *A. mexicanus*^38,40^ that have evolved independent of eye degeneration. Gross et al.^41^, however, found a correlation of pigment loss with blindness in cave fish, suggesting it might also be driven by selection. Yoshizawa et al.^42^ recently demonstrated that neural crest cells are key agents in cavefish eye and pigment regression, as transplanting embryonic neural crest cells from surface-into cave-dwelling fish embryos resulted in increased eye size and induced pigmentation and growth of melanophores in the cave fish. This suggests the possibility of common underlying factors for eye degeneration and pigmentation loss.

In addition to the observed regressive traits, we found evidence that cave loach developed constructive traits that compensate for the loss of vision. Cave loach and hybrids had significantly longer barbels and larger olfactory epithelium than measured in surface fish, regardless of treatment. Elongated appendages including barbels are often found in cave animals including fish^43^ and are typically associated with enhanced tactile and chemosensory abilities. More extensive olfactory epithelium will probably increase identification of chemical cues associated with food, communication, and reproduction. In some fish with an exceptionally sensitive olfactory sense, including sharks, the size of the olfactory epithelium can be correlated with olfactory^44,45^. However Blin et al.^46^ reported that the enlarged olfactory epithelium does not account for the enhanced amino acid detection skill of cave forms of *A. mexicanus*. Our findings suggest that mechano- and chemo-sensory systems are vital in darkness and, hence, expanded in cave loach. Laboratory trials might elucidate the survival potential of such effects. Studies of cave *A. mexicanus* have found increased numbers of taste buds and enhanced chemosensory and food detection capabilities^2,10^. Future studies of cave fish, including our study species, will demonstrate the extent of universality of such adaptations.

It is clear from the differences among the cave, hybrid, and surface loach across the two light conditions that targeted traits must have a heritable basis. However, we also observed phenotypic plastic responses to light conditions for all loach studied, indicating a gene/environment interaction. Offspring of cave fish that developed from the fertilized egg to the juvenile stage (∼ two months) in the natural photoperiod developed 14% larger eyes, 19% darker body coloration, 18% shorter barbels, and 8% smaller olfactory epithelium compared to their siblings raised in total darkness. The phenotypic responses were present in the first generation. This phenomenon has been described for several troglomorphic animals including fish and crayfish^47^. These results strengthen the suggestion that phenotypic plasticity, probably jointly with epigenetic regulator activity^30,48^, might play a fundamental role in the adaptive processes of cave fish^49^. The ability to quickly adapt to environmental conditions is essential to all organism survival. Phenotypic plasticity plays a major role, allowing individuals to initially survive an abrupt change of environment (as in being trapped underground), with subsequent selection triggering troglomorphic adaptation. In this way phenotypic plasticity could, in the long run, pave the way for genetic adaptations^48,50,51^.

## Conclusions

Cave organisms provide excellent models for the study of evolutionary principles. Cave loach bred in captivity develop typical troglomorphic adaptations including regressive traits, the reduction of eyes and pigmentation, and constructive traits, elongation of barbels and increased size of the olfactory epithelium. The morphology of this cave fish is affected by a combination of heritable genetic differentiation and phenotypic plasticity. The eye lens deterioration observed in F1 cave loach but not in hybrids suggests deleterious changes in the genes or gene functions, while the high variation in eye size and pigmentation observed among cave fish may reflect its genetic history or ongoing introgression between surface and cave fish. Phenotypic plasticity observed in response to light conditions highlights the ability to quickly adapt to a novel environment. The ability to keep and breed cave loach in the laboratory and create hybrids of cave and surface populations, highlights this newly discovered representative of troglobiotic fishes as a unique opportunity for further work on evolutionary adaptation including physiology, behaviour, genetics, and ecology.

As most of the work on troglomorphic adaptation in fishes has been done with the Mexican tetra, comparative studies with cave loach can potentially elucidate species-specific developmental and genetic pathways and characterize convergent phenotypic changes such as in eye and pigment degeneration.

## Acknowledgements

We are most grateful to cave diver Joachim Kreiselmaier. His tenacious and enthusiastic participation, sacrificing much of his free time to dive in cold subterranean waters, and his skill in capturing cave loach under extremely challenging conditions enabled this study. We are grateful to all members of the “Freunde der Aachhöhle e.V” for their kind support. Warm tanks to family Bürßner for allowing entry onto their property to access the entrance to the Danube-Aach sytem. We thank Myriam Schmidt for her help in fish maintenance. This work was financially supported by the University of Konstanz to J.B-G and A.Bö. We thank Lucidus consultancy for English language editing.

## Notes

### Competing Interest Statement

The authors have declared no competing interest.

