## Supplemental information for "Phenotypic plasticity and genetic differentiation drive troglomorphic character development in European cave loach"

**Supplementary figures**

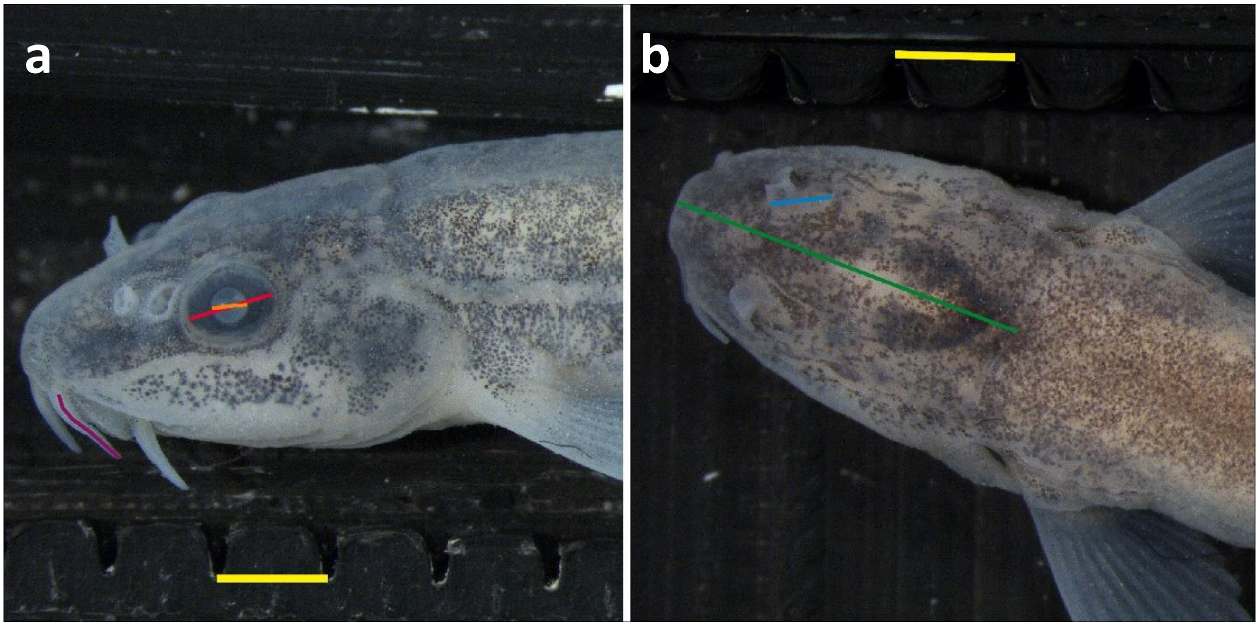


**Fig. S1. (a)** F1 offspring of surface loach *Barbatula barbatula* Scale bar (yellow line) = 2 mm. red line = eye diameter; orange line = lens diameter; purple line = second barbel length. **(b)** Dorsal view F1 offspring of cave loach reared in complete darkness. Eyes are retracted into head. Scale bar (yellow line) = 2 mm. blue line = olfactory epithelium diameter; green line = head length.

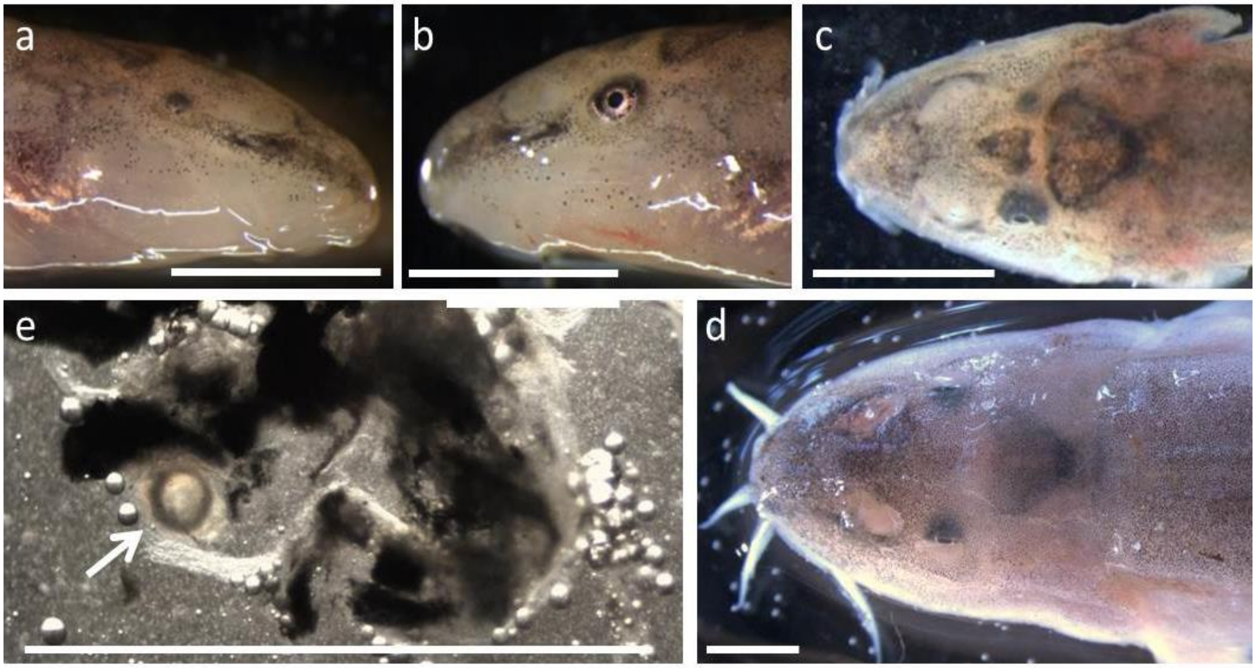


**Fig. S2. Eye regression in wild cave loach sampled in the Danube-Aach system. (a–c)** Bilateral asymmetry in eye development of a single fish; **a, c**: rudimentary eye on right side, **b, c**: reduced but visible eye on left side. **(d, e)** cave loach with small retracted eyes, **d:** dorsal view of head, **e:** squash preparation of both eyes showing one small lens only (arrow). Scale bar = 3 mm

### Supplementary methods

*<u>Access to the Danube Aach system and specimen collection</u>*

The Danube-Aach cave system was accessible to divers only in dry summer months of low water flow (Fig. 1b). The system is complex with multiple narrow passages. The bottom is mostly covered with silt, creating turbidity. Visibility averages 1–2 m (J. Kraiselmaier, personal observation). Divers were forced avoid diving near the bottom, the usual habitat of loach, possibly the reason that the cave fish was not detected sooner, as the system has been explored since the early 1980s. The only entry point besides diving from Aach spring, is via a shaft dug out within 13 years resulting 29 November 2003 with the discovery of Doline Lake, ~100 m underground (Fig. 1b). Since 2003 this has allowed exploration of the Danube-Aach cave system further upstream toward the Danube even beyond a collapse zone. Cave fish have not been detected in the small air-filled cave area that can be reached on foot at Doline Lake (Fig.1b).

Each diving trip lasted a minimum three hours including decompression time (maximum water depth ~ 30 m). The success rate of cave loach capture was low. However, loach were frequently observed during the 21 dives totalling 57 hours from 2015 through 2018. The diver described fish of all age classes. On two dives in September 2017 and summer 2022, diver observed more than 50 young of the year cave loach, some of which were caught in 2017, confirming that the cave loach regularly reproduces in the Danube-Aach cave system.

*Breeding of surface fish from Danube and Radolfzeller Aach*

Late January 2019 when water temperature was ~ 7.5°C, we selected four loach pairs from Danube and four from Radolfzeller Aach laboratory populations (Table 1). One male and one female were hand netted and transferred to a circular 20 l tank with water flow-through from Lake Constance and an air stone for aeration. The light regime followed the natural photoperiod. At the beginning of March, the water temperature, which had reached ~12°C, was slowly raised to 14°C over two days. We introduced a removable plastic sheet covering the tank bottom for egg collection. The tanks were checked for eggs each day and if detected, the plastic sheet was carefully removed and cut in half for transfer of the adhering fertilized eggs to the common garden experimental setting, and a new plastic sheet was placed in the tank.

### *Crossbreeding*

In the same room and at the same time as the breeding of the surface fish, three pairs were separated for the production of hybrids (Table 1). Two pairs included a male from Radolfzeller Aach and a female cave fish. One pair included a male from the Danube and a female cave fish. We lacked a sufficient number of captured cave fish males to include a pair with a cave fish male and surface fish female. Treatment was identical to the production of surface fish offspring.

### *Breeding of cave fish from Danube Aach system*

At the end of January 2019, cave fish breeding was initiated. Four pairs of male and female cave fish were placed in individual 60 l tanks (Table 1). All fish except one male were fish captured from the wild. One male was an F1 offspring from a breeding event that took place in 2018 in the captured cave fish population. Multiple Petri dishes containing stones were added to the tank as substrate for eggs, at a later date we switched to covering the bottom with a removable plastic sheet as described above. To induce breeding, water temperature was gradually raised to 14°C. Due to failure of breeding, an additional cave fish pair (family C6; Table 1) was isolated on 15 May 2019. This was necessary to produce more cave fish offspring for the common garden experiments.

### *Maintenance and breeding of captured loach*

Cave and surface loach consumed a range of food items including blood worms, daphnids, and several types of artificial food (Tetra flakes, trout pellets). With increasing water temperature in spring, surface loaches could be observed spawning, primarily shortly after sunrise. When spawning, both sexes swam upward in the water column in pairs, releasing eggs and sperm simultaneously in open water. Eggs sank and adhered to the substrate. Loach produced a high number of offspring and exhibited no brooding care behaviour. Neither eggs nor offspring were consumed by adult loach, and there was no need to remove the fertilised eggs or hatched fry from the maintenance tanks. Both Danube and Radolfzeller Aach surface populations successfully reproduced during spring 2017 and 2018.

In April/May 2018, two male and two female adult cave loach (the only adult cave fish available by that time), were placed together in a 60 l tank with constant water flow from Lake Constance. The water was gradually warmed from 8°C to 15°C over the course of four weeks. Spawning substrate, consisting of sand, fine and coarse gravel, and flat stones was provided. In July 2018, four juvenile cave fish of ~2 cm were detected in the tank. These fish represented the first successful breeding of cave loaches from the Danube-Aach cave system in captivity. Although we checked, no eggs were seen in the tank during the short time that the light was switched on for daily feeding or cleaning. We presume that most offspring were lost due to starvation, since no special young fish food had been provided before the detection of juvenile fish.
